# Creating a Community of Data Champions

**DOI:** 10.1101/104661

**Authors:** Rosie Higman, Marta Teperek, Danny Kingsley

**Affiliations:** Office of Scholarly Communication University of Cambridge

## Abstract

Research Data Management (RDM) presents an unusual challenge for service providers in Higher Education. There is increased awareness of the need for training in this area but the nature of discipline-specific practices involved make it difficult to provide training across a multi-disciplinary organisation. Whilst most UK universities now have a research data team of some description, they are often small and rarely have the resources necessary to provide targeted training to the different disciplines and research career stages that they are increasingly expected to support.

This practice paper describes the approach taken at the University of Cambridge to address this problem by creating a community of Data Champions. This collaborative initiative, working with researchers to provide training and advocacy for good RDM practice, allows for more discipline-specific training to be given, researchers to be credited for their expertise and an opportunity for those interested in RDM to exchange knowledge with others. The ‘community of practice’ model has been used in many sectors, including Higher Education, to facilitate collaboration across organisational units and this initiative will adopt some of the same principles to improve communication across a decentralised institution. The Data Champions initiative at Cambridge was launched in September 2016 and this paper reports on the early months, plans for building the community in the future and the possible risks associated with this approach to providing RDM services.

## Introduction

Whilst funder and institutional policies have introduced drivers for Research Data Management (RDM) there is still much work to be done to engage the majority of researchers. There is a need to raise their awareness of appropriate RDM and to explain the benefits of sharing their research data. This is proving to be a labour-intensive and lengthy process due to the wide variety of forms RDM can take between and within different disciplines. The Research Data Facility at the University of Cambridge has been successful in engaging with academic staff. The training programme is heavily subscribed but currently lacks stable funding to employ enough people to meet this demand. Aside from the resourcing issues it is also difficult for any central support service to develop the expertise needed to provide in-depth advice in every research-active discipline, from Architecture to Zoology, given the range of data and research methods that these entail.

As well as training researchers, the Research Data Facility offers individual advice on data management plans, support to deposit data in the institutional repository, and consultancy on all aspects of RDM. The results of the recent data asset framework survey conducted with Jisc (Johnson, Chiarelli, & Parsons, 2016) revealed that 46% of researchers at Cambridge who were not aware of the support services available to them. This survey was self-selecting, in that the people who responded are already engaged enough to participate. This finding underlines the challenge ahead for staff providing RDM services; the Research Data Facility at Cambridge had conducted 99 departmental briefings in the 18 months prior to the survey.

These departmental briefings to researchers were the first form of advocacy by the Research Data Facility and took the form of information sessions focused on compliance and so failed to engage researchers (Teperek, Higman & Kingsley, forthcoming). We have subsequently revised our approach to focus on the underlying reasons for good RDM and how this can benefit researchers, which has improved feedback and researchers’ attitudes towards RDM. However, information sessions can only provide an overview and cannot be targeted to different career stages when researchers will typically be motivated by different pressures and incentives. To allow for more targeted advocacy the briefings are now supplemented by training sessions aimed at PhDs and postdoctoral researchers, and are tailored for specific disciplines, something previous research has highlighted as important for engagement (Hiom, Fripp, Gray & Snow, 2015). These training sessions allow for in-depth discussions of the wider issues of back-up, managing data, and researching in an open manner in an interactive workshop (Teperek & Higman, 2017a). They have been extremely successful but are labour intensive to prepare and deliver, and so it is impossible to meet demand within currently available resources.

Faced with the dual problems of insufficient resources to meet the demand for training and the need to raise awareness of our research data support services, and RDM more generally, we have set out to create a community of Data Champions across the University. Data Champions are researchers, PhD students or support staff who have agreed to advocate for good data management and sharing practice within their department: providing local training, briefing staff members at departmental meetings, and raising awareness of the need for data sharing and management. This paper will outline the community-focused approach taken, activities of the Champions thus far, the risks associated with the project and our future plans to develop this community.

## Building on a Collaborative Approach

The Data Champions initiative builds on the existing approach to RDM support services at Cambridge, emphasising the benefits of data management and sharing ahead of compliance and aiming to work in a collaborative fashion (Teperek, Higman & Kingsley, forthcoming). Recognising that RDM needs to be a researcher-owned endeavour to ensure its sustainability, we actively engage with the researcher community. This includes being part of the OpenCon Cambridge satellite group^1^ which is largely comprised of early career researchers. This has allowed us to build relationships with individuals who are passionate and willing to dedicate time to advocating for openness. The Research Data Facility also collaborates with a project group made up of support professionals and researchers who work to improve our RDM services. Collaboration with researchers and other staff is fundamental to achieving the cultural change necessary for the normalisation of data sharing and management.

Taking a similar approach, we have attempted to engage with a wide variety of groups in setting up the Data Champions initiative. We used two parallel approaches to reach out to our target audiences: sending information via a range of mailing lists and social media to all research and research support staff, and a targeted approach to engage individuals with specific expertise we thought it would be useful to share across the University. The initial advert described what is required of the Data Champions as well as outlining how this will benefit them and their careers (Higman and Teperek, 2017b). There is a Data Champions website^2^ ensuring that individuals receive credit and recognition for the time and expertise that they contribute, as well as opportunities for networking, additional training and developing leadership skills. There were minimal requirements in the advert, emphasising an enthusiasm for data sharing and management, to engage the broadest possible group of researchers and support professionals.

This advert resulted in 43 eligible applicants from Cambridge who we approached about attending an initial meeting for the Data Champions (Higman and Teperek, 2017b). The vast majority of applicants came from Science, Technology, Engineering and Medicine (STEM) subjects, reflecting the general pattern of engagement we have witnessed with RDM support services. Researchers made up 72% of applicants, and the fact that the programme is researcher-led ensures sufficient domain knowledge to be able to effectively tailor the template workshops for individual departments. The aim of the programme is that other researchers engage with the Champions as peers in their discipline. The largest group of applicants were PhD students and postdoctoral researchers, making up 24% and 36% respectively, but there are also several lecturers and principal investigators involved who we hope will be able to provide leadership to the more junior participants. We decided to accept all eligible applicants, meaning that there are several Champions in some departments, to encourage the enthusiasm that applicants had displayed and in recognition of the likely attrition there will be as the project progresses.

## Establishing a Community of Practice

Building on the collaborative approach used for recruiting the Data Champions we would like them to become a ‘community of practice’ (CoP) (Wenger, 1998) as well as a means to increase the quantity and diversity of training being provided. There are many passionate individuals involved who have extensive knowledge and skills in different aspects of data management; a CoP would be a good way of achieving the aim of increasing RDM knowledge exchange across university. Knowledge exchange is particularly important as the University of Cambridge is a highly decentralised institution spread across a city with numerous units who are quite independent of each other. Furthermore, much of this RDM knowledge is tacit, used in day to day practice but not well documented, so hard to capture without relationships and regular interactions between key individuals (Wenger, McDermott, & Snyder, 2002).

A CoP is defined as “groups of people who share a concern, a set of problems, or a passion about a topic, and who deepen their knowledge and expertise in this area by interacting on an ongoing basis” (Wenger, McDermott & Snyder, 2002). Whilst the Data Champions are clearly not a CoP in the strictest sense as some structure and demands are being imposed centrally, we hope to be able to take many of the principles of a CoP and foster a sense of shared purpose in the group. This will also fit with the researcher-led approach to RDM we are trying to take elsewhere in our services. There is a delicate balancing act required when facilitating a CoP instigated by an organisation rather than individual participants. Organisational support can lend the group greater legitimacy, something we are hoping to provide by writing to each Head of Department informing them of the local Data Champion and that they have our support, but there is also a risk of over-management (Wenger, McDermott & Snyder, 2002). This is a particular risk in this project as we have set the initial agenda for the group which may affect later adopters (Hodgkinson-Williams, Slay & Siborger, 2008). To combat these risks we have minimised the demands placed on the group and are seeking regular feedback and input from the Champions.

Notwithstanding these risks, there are already some aspects of a CoP in the setup of the Data Champions: participation will help individuals develop their skills while also contributing to the wider organisational goal of promoting better RDM (Wenger, McDermott, & Snyder, 2002). Based on the Champions’ initial applications the opportunity to develop their knowledge of RDM motivates some whilst others are driven by promoting open research and data sharing and how these impact on their research. These dual purposes provide a domain for the CoP to focus on and practices which they can develop together which will be important given the diverse nature of the group. One advantage of launching this type of initiative in academia is that it will align with many researchers’ existing experiences of being in an, often implicit, CoP based around their discipline and location (Nistor, Daxecker, Stanciu & Diekamp, 2015). We are hopeful that with organisational support provided by our team, flexibility in how they want to engage, and a clear domain and practices to develop determined by the Data Champions’ interests, this group can be an active CoP as well as training their colleagues.

### Welcoming our Data Champions

Turning the initial expression of enthusiasm into a functioning CoP is crucial to the long-term success of the project, and to this end we have organised a series of events to welcome the Data Champions and help them feel part of a community. Initial welcome meetings were held in December 2016, across two sites to make them as accessible as possible, and were attended by 33 Champions and supporting librarians. The meetings introduced Champions to our expectations from them (whilst giving space for concerns and alternative ideas for activities to be raised), the support available to them and presented the template workshop for STEM subjects which Champions will be asked to tailor for their own departments (Higman and Teperek, 2017a). We also demonstrated our prototype Data Champions website with profiles pre-populated for the support staff involved to encourage the Champions to create their own profile^3^. In the future we hope to develop this website further so that users can search for Champions by department and areas of expertise.

The welcome meetings were kept deliberately informal with a break for lunch, and time for discussions and networking. Allocating time for building relationships was prioritised as these connections can give an incentive for people to participate and feel comfortable doing so, especially as many did not know one another and sit in different parts of a large organisation (Eberle, Stegmann & Fischer, 2014; Hodgkinson-Williams, Slay, & Siebörger, 2008). The Data Champions suggested many ideas at the meetings, leading to conversations in person, over email and on Twitter (#datachampcam) in the following weeks that helped to build a sense of community and purpose.

Whilst there was much enthusiasm evident it was also clear that there is a wide variety in the Champions’ confidence and willingness to engage. Some expressed doubt about their ability to deliver one training session a year (the minimum requirement we set) whilst others felt that this was far too little and are planning more ambitious advocacy in their departments. One training session per year was set as a minimum requirement for each Champion in response to the Research Data Facility’s lack of resources to deliver training – the initial trigger for the initiative. However, recognising that this will be intimidating for some, we have also suggested smaller scale advocacy to help build the Champions’ confidence, such as a briefing session for their research group. This flexibility is important given the range of participants we have, and we hope that providing easier tasks for the less experienced will make them feel welcome in the group (Eberle, Stegmann, & Fischer, 2014).

Some Data Champions could not attend the welcome meetings so the materials were made available to everyone afterwards and we met several Champions individually or in small groups to ensure that they felt part of the community and understood the support available and expectations of them. Although it is still too early to tell whether the group will coalesce into a sustainable community, there are already similar patterns of engagement as seen in CoPs. There are relatively small ‘core’ and ‘active’ groups at the centre who are driving activities within the group and their departments, and a larger periphery who are contributing occasionally and learning through observation (Wenger, McDermott, & Snyder, 2002).

As well as in-person welcome meetings, a space was created for the Data Champions in the institutional virtual learning environment (VLE) where presentations can be shared, news posted, and there is a forum for discussion. We have tried to seed activity in this space by encouraging those who already deliver RDM-related sessions to upload their presentations, but otherwise there has not been a great deal of activity here. There are several possible reasons for this including poor design of the site, fears of being the first one to post, and participants having been away for much of the time it has been live due to the Christmas break. We are reviewing the design of the space in order to combat the first problem and will be seeding discussions on the forum by encouraging those who email us directly to make a public post instead where appropriate. If these measures are unsuccessful we will investigate other communication tools, such as Slack, to see if these are more conducive to discussion.

In addition to the VLE site there is a Data Champions email list which is used to advertise relevant events and opportunities and upcoming training and meetings for Data Champions. We have also created a Twitter hashtag (#datachampcam) for the group which is gradually being adopted and is useful as Twitter is a space already used by many of our Champions. It is quite hard to determine how useful these electronic spaces are as they may be frequented by ‘lurkers’ who are not confident enough to offer their own views but are still learning. Despite the current low levels of engagement within the VLE it remains a useful space for sharing materials with the Data Champions and we will be working on encouraging more activity there in the coming months.

### Developing Data Champions

The period immediately following the first meeting has been identified as a particularly vulnerable time in CoPs as the initial energy can dissipate and participants may not yet have formed the relationships which tie them to the group (Wenger, McDermott, & Snyder, 2002). In recognition of this we have invited the Data Champions to two events being run by the Research Data Facility and run presentation skills training for those who lack confidence presenting to their peers. The presentation skills training formed part of our offer to train up Data Champions and support the less experienced members of the community so that they feel more comfortable advocating to their more senior peers. These entirely optional sessions were quite well-attended with 12 people coming over two sessions, with good feedback from participants. The training was an adaptation of a session run regularly by the Office of Scholarly Communication for librarians, with more time allowed for practising our template RDM presentation^4^. The presentation skills training also provided a useful opportunity for Champions to ‘check-in’ in person, discuss any concerns that they may have about being a Data Champion and share ideas about activities that they are planning. This facilitated the Champions in continuing to build relationships within the group and maintain the momentum from the welcome meetings.

In the future these activities will largely take place at the bi-monthly ‘forums’ where the Data Champions can report back on their activities and share their expertise. These will also help to establish a regular ‘rhythm’ to the community, where people know that they will see one another, and so help keep Data Champions engaged (Wenger, McDermott & Snyder, 2002). The first bi-monthly forum meeting will be held in March 2017 over a two hour lunch with several short presentations by Data Champions on aspects of RDM, an update from the central Research Data team on services and events, and a facilitated discussion on what does and doesn’t work when advocating. In the spirit of a collaborative CoP we will solicit feedback at the end of this meeting to find out if the format meets Champions’ expectations, and are considering a more social event over the summer to help build relationships in our community.

In addition to these centrally organised events the Data Champions have also begun to advocate in their departments. The activities so far include:

- A training needs analysis and FAQs developed^5^ in Chemistry by the librarian and the Data Champions;
- A sub-group being formed in Engineering where there are several champions and an active library service;
- Advertising via departmental newsletters;
- Briefing talks to academic staff from several Data Champions.

Planned activities include regular short tips about RDM via email and lunchtime talks that fit into an existing departmental series. Several Champions are also approaching their local IT officers as IT services have implications for the approaches which can be suggested by the Data Champions. The variety of approaches being taken reflect the different roles of Data Champions and existing methods of communication with departments. This is encouraging as it reflects our supposition at the beginning of the project that we were not in a position to understand the variety of ways in which each department communicates. In keeping with attempts to make this a community-driven initiative the Champions have been encouraged to consider their role and department, and then advocate in the way they feel will be most effective, with the proviso that at least one teaching session is delivered per year.

## Risks of the project

Whilst our initial call for participants and the meetings held so far have shown that there is interest among Cambridge researchers and staff in developing a network of Data Champions, there are clear risks to the project. These can be classified into three main problems:

- Insufficient rewards for participating
- Champions participating but not delivering training
- Insufficient resourcing to support the group

### Insufficient rewards for participating

A major risk of the project is that Champions will not feel sufficiently rewarded for the time that they are volunteering to the group and so either not actively participate or leave. This is a difficult risk to manage as we do not have the resource to support any formal reward system. Instead we are focusing on providing them with publicity, networking and training. We have written to the Head of Department for each Champion to tell them that their staff have volunteered and have our backing. We promote the Data Champions via our Twitter feed and in our presentations where appropriate. They are encouraged to create a profile on our website where they can advertise their expertise^6^. We will also be providing them with a graphic for their email signatures so that they can easily advertise their new role to their colleagues. The Data Champions are also provided with access to training, such as the presentation skills sessions which would not otherwise be available, and the opportunity to talk to others with similar interests and problems. Some of our Champions are likely to participate regardless of the rewards on offer as they are passionate advocates of Open Research, but many need to see some benefits from participating so unless we get the rewards right there is a risk that we will not expand the network of people talking about RDM in the University.

### Champions participating but not delivering training

The Data Champions community is made up of people with wide-ranging roles including PhD students, lecturers, data managers and librarians. These groups are likely to participate in different ways but based on activities so far there is a risk of many people participating actively but not delivering the training sessions that were the initial driver for the initiative. Some of the senior researchers are participating actively but are more interested in delivering short briefing sessions than full workshops. Others are particularly interested in promoting data sharing rather than RDM more generally. The less experienced members of the group are interested in learning about RDM at the moment so will want to take on a peripheral role where they mostly observe. This creates a dilemma as the advocacy being proposed could be extremely useful in promoting RDM but will not necessarily satisfy the need for training which the project was intended to meet. This risk will need to be monitored to ensure that the activities taking place justify the resource being dedicated to the project.

### Insufficient resourcing to support the group

Without most of the Data Champions delivering workshops or briefing sessions it would be hard to justify the resources being put into supporting the Data Champions. Inevitably a large community such as this takes time to set up and run; from the formal events, to speaking with individuals and connecting people with similar interests and needs. It is estimated that running a CoP will take 20-50% of the community coordinator’s time (Wenger, McDermott and Snyder, 2002), something which is not budgeted for in our current staff allocation. This can be justified if there is a large scale increase in the amount of advocacy for RDM, although it may still end up taking more of the Research Data Facility’s time in total. Therefore, the priority for the coming months is to make sure that the training happens and consider how we can best demonstrate the value of the project.

## Demonstrating value

Given the time invested in this project by the Research Data Facility, departmental librarians, and Data Champions, it is crucial that we can demonstrate the value it produces. This can be hard in any CoP and particularly so when, as with the Data Champions, much of the impact will be in small, intangible changes in practice and attitudes. Furthermore it will be difficult to isolate the impact of the Data Champions from that of the Research Data Facility more generally. For example, we have recently upgraded the system for submitting research data into the institutional repository, so if there is an increase in submissions to our repository it will be hard to determine whether it was due to the upgrade or the advocacy of our Data Champions.

We are still deciding what measures of value to adopt in the long-run but are initially collecting data on how many people have attended sessions run by Data Champions and what proportion of them felt that it was beneficial. Although this will not capture individual conversations Data Champions may be having it will allow us to see the scale of activity from the group and should highlight any major problems with the sessions being run. This should mitigate a further risk with this project which is that some of the training being delivered may not be of the quality expected. As well as asking the Data Champions to collect data on whether the sessions were perceived to be useful, we will also provide a template feedback form in case they wish to collect more detailed feedback (Higman and Teperek, 2017b). We are considering more extensive data collection but are also wary of the risk of imposing too many administrative burdens on people who are volunteering their time for relatively few tangible rewards.

## Maintaining our Community of Data Champions

The initial expression of interest from our research and support staff community was encouraging and has been converted into a nascent community who are building relationships and starting to take action. To cultivate this CoP and ensure its sustainability in the long-run will require a stable base of members and a more well-defined domain for the group to focus on.

To help maintain the size of the group we have asked Champions to identify someone appropriate in their department or research area who could replace them if they plan to leave. We are also advertising the Data Champions in information sessions and workshops which we run centrally to help garner interest in group, especially in subject areas where we do not currently have a Champion. This will hopefully outweigh the inevitable attrition as people’s interests and jobs change so they have to leave the group. Whilst we are taking these steps to grow the CoP the focus for now will be on building relationships within the group to ensure people remain involved, rather than building a huge membership base (Wenger, McDermott & Snyder, 2002). Research into other CoPs in academia have demonstrated the importance of spending time in communities and building relationships in order to develop a sense of community (Nistor, Daxecker, Stanciu, & Diekamp, 2015). This will form the basis for most of our activity in the next six months.

In the slightly longer term it will be important to better define the ‘domain’, the issues on which the Data Champions will focus. Currently the focus of the group reflects the areas the Research Data Facility supports and the interests of the individuals involved, and has not been explicitly defined. This is appropriate in the initial stages so the community can accommodate different interests but at some point the group will need to set boundaries to define its scope (Wenger, McDermott & Snyder, 2002). This is particularly important as the Research Data Facility are looking at new areas such as software and electronic lab notebooks and this raises questions over whether the Data Champions can be expected to cover these areas as well.

## Conclusion

Based on our experience so far, a Data Champions initiative seems to be an effective way to increase both advocacy for RDM and discipline-specific training available to researchers in large universities. A CoP around RDM allows existing experts and interested individuals to exchange knowledge and develop ways of influencing their colleagues. Our experience suggests that launching this type of initiative after a period of engaging with researchers produces a positive response. Instigating a CoP from a central support service runs a risk of over-management and researchers not engaging so it will be important that we take a sufficiently community-driven approach in all our future activities with the Data Champions.

The community we are building with the Data Champions should address some of the data and domain skills gaps in RDM training, but there are many issues which it clearly cannot address. The incentives structure in higher education persists in placing undue emphasis on publishing in high impact journals. Such issues cannot be touched by the Data Champions initiative^7^, requiring alternative approaches such as lobbying at a senior level. However we hope that the Data Champions initiative will build a community of engaged researchers, who are changing the culture within their own disciplines.

## Acknowledgements

Many thanks to the RDM Outreach and Support group: Clair Castle, Jennifer Copic, Georgina Cronin, Jenifer Wright for their help in getting the project started, ongoing input and invaluable advice.

## Footnotes

1 OpenCon Cambridge: http://www.openconcam.org/

2 Data Champions: http://www.data.cam.ac.uk/intro-data-champions

3 Data Champions: http://www.data.cam.ac.uk/datachampions

4 Presentation skills for Data Champions: http://www.slideshare.net/ClaireSewell/presentation-skills-fordata-champions

5 Open data FAQs for chemists: http://www-library.ch.cam.ac.uk/open-data-faqs-chemists

6 Data Champions: http://www.data.cam.ac.uk/datachampions

7 Jisc Research Data and related topics: https://researchdata.jiscinvolve.org/wp/2016/09/27/research-datamagic-anyone

